# methylscaper: an R/Shiny app for joint visualization of DNA methylation and nucleosome occupancy in single-molecule and single-cell data

**DOI:** 10.1101/2020.11.13.382465

**Authors:** Parker Knight, Marie-Pierre L. Gauthier, Carolina E. Pardo, Russell P. Darst, Alberto Riva, Michael P. Kladde, Rhonda Bacher

## Abstract

Differential DNA methylation and chromatin accessibility are associated with disease development, particularly cancer. Methods that allow profiling of these epigenetic mechanisms in the same reaction and at the single-molecule or single-cell level continue to emerge. However, a challenge lies in jointly visualizing and analyzing the heterogeneous nature of the data and extracting regulatory insight. Here, we developed methylscaper, a visualization framework for simultaneous analysis of DNA methylation and chromatin landscapes. Methylscaper implements a weighted principle component analysis that orders sequencing reads, each providing a record of the chromatin state of one epiallele, and reveals patterns of nucleosome positioning, transcription factor occupancy, and DNA methylation. We demonstrate methylscaper’s utility on a long-read, single-molecule methyltransferase accessibility protocol for individual templates (MAPit) dataset and a single-cell nucleosome, methylation, and transcription sequencing (scNMT-seq) dataset. In comparison to other procedures, methylscaper is able to readily identify chromatin features that are biologically relevant to transcriptional status while scaling to larger datasets.

**Availability and implementation:** Methylscaper, is available on GitHub at https://github.com/rhondabacher/methylscaper.

## Introduction

Abnormal epigenetic changes are a key hallmark of cancer. Alterations in DNA methylation, including the co-occurrence of both hyper- and hypo-methylation of different regions of the genome, have been detected in nearly all cancer types(1–3). Additionally, both cancer- and tissue-specific differences exist in nucleosome positioning and occupancy, as well as transcription factor binding activity, which determine chromatin accessibility (4). However, profiling endogenous methylation and accessibility states separately ignores their complementary nature in regulating gene expression and, by definition, queries different sets of molecules (5). To address this, assays such as MAPit-BGS(6) and NOMe-seq(7) that simultaneously capture nucleosome occupancy and methylation states at single-molecule resolution have been developed. In both cases, chromatin accessibility is first probed by the methyltransferase M.CviPI(8), which methylates unprotected GC sites. Next, accessibility at GC sites and CG endogenous methylation are profiled by bisulfite(9) or bisulfite-free enzymatic conversion(10). After sequencing, the methylation signals of all cytosines are translated bioinformatically. Long-read sequencing is particularly advantageous to phase the co-occurrence of epigenetic features, e.g., multiple nucleosomes. Recently, an extension of NOMe-seq, nanoNOMe, made use of long-read nanopore sequencing and resolved long-range patterns along individual DNA molecules(11). Methods for simultaneously profiling accessibility and methylation have also been extended to single cells via the scNOMe-seq(12) and scNMT-seq(13) techniques.

For MAPit and nanoNOMe, the long reads derive from contiguous single DNA molecules, while single-cell methods use short reads that are reconstructed into contiguous DNA molecules from individual cells. Both types of methods allow for discerning the heterogeneous nature of cellular DNA methylation and chromatin structure. Bioinformatic software programs, such as Bismark(14), are used to align the data; however, many analytical pipelines and downstream visualization tools fail to highlight the epigenetic variation in a useful way. Previously developed methods utilize the output from Bismark but are limited to a relatively small number of reads or provide summary plots rather than site-level data(15,16). Two other such visualization tools are the NOMePlot(17) and MethylViewer(18) applications, which were designed to simultaneously visualize CG methylation/GC accessibility patterns. Despite their integrated pipelines, the commonly used ‘lollipop’ plots are not intuitive in highlighting the joint occupancy and methylation states along a continuous DNA strand, especially when considering hundreds or thousands of molecules. The previously developed MethylTracker(19) plots visually intuitive methylation/accessibility patterns by connecting consecutively methylated or unmethylated sites with contrasting colors, however it is computationally inefficient and unable to effectively organize hundreds of reads.

Here, we describe methylscaper, a bioinformatic and statistical software package that processes raw sequencing reads and generates visualizations of the DNA methylation and chromatin accessibility patterns. Additionally, output from Bismark for single-cell joint profiling experiments may also be used as input. Ordering the reads is a key step for visualization and our pipeline implements a two-stage weighted principal component analysis (PCA) framework that is feature and site specific. Weighting allows the user to emphasize specific genomic regions or features of interest. Compared to alternative procedures, our ordering is also efficient for large-scale datasets. Methylscaper is an interactive visualization platform available as an R/Shiny application and may also be used directly via the R package. We evaluate methylscaper on an epigenetic DNA resilencing MAPit-BGS dataset and demonstrate its superior ability to elucidate epigenetic patterns. We further demonstrate methylscaper on a single-cell dataset generated using scNMT-seq and identify regions of cell-to-cell nucleosome sliding.

## Methods

Methylscaper first performs pre-processing to align the raw sequence files, followed by visualization and statistical analysis of methylated and accessible chromatin regions. The initial pre-processing steps include pairwise alignment of each sequence, quality control and filtering of poorly aligned sequences, and finally, conversion of the aligned sequences to methylation and occupancy states (Figure 1A). Additional details on the bioinformatic processing are available in Supplementary Materials. Regions of methylation or accessibility are identified by connecting consecutive sites having the same methylation state (Figure 2B). A patch of endogenous methylation is plotted in red if ≥2 consecutive HCG sites show methylation (C in the sequence). Similarly, consecutive GCH (H=A,T, or C; details in Supplementary Materials) methylation indicates accessibility, plotted in yellow. By contrast, consecutively unmethylated GCH or HCG (T in the sequence) are colored black. Patches of any color are interrupted by single GCH or HCG of the opposite methylation state, which are emphasized as gray borders.

**Figure 1:**
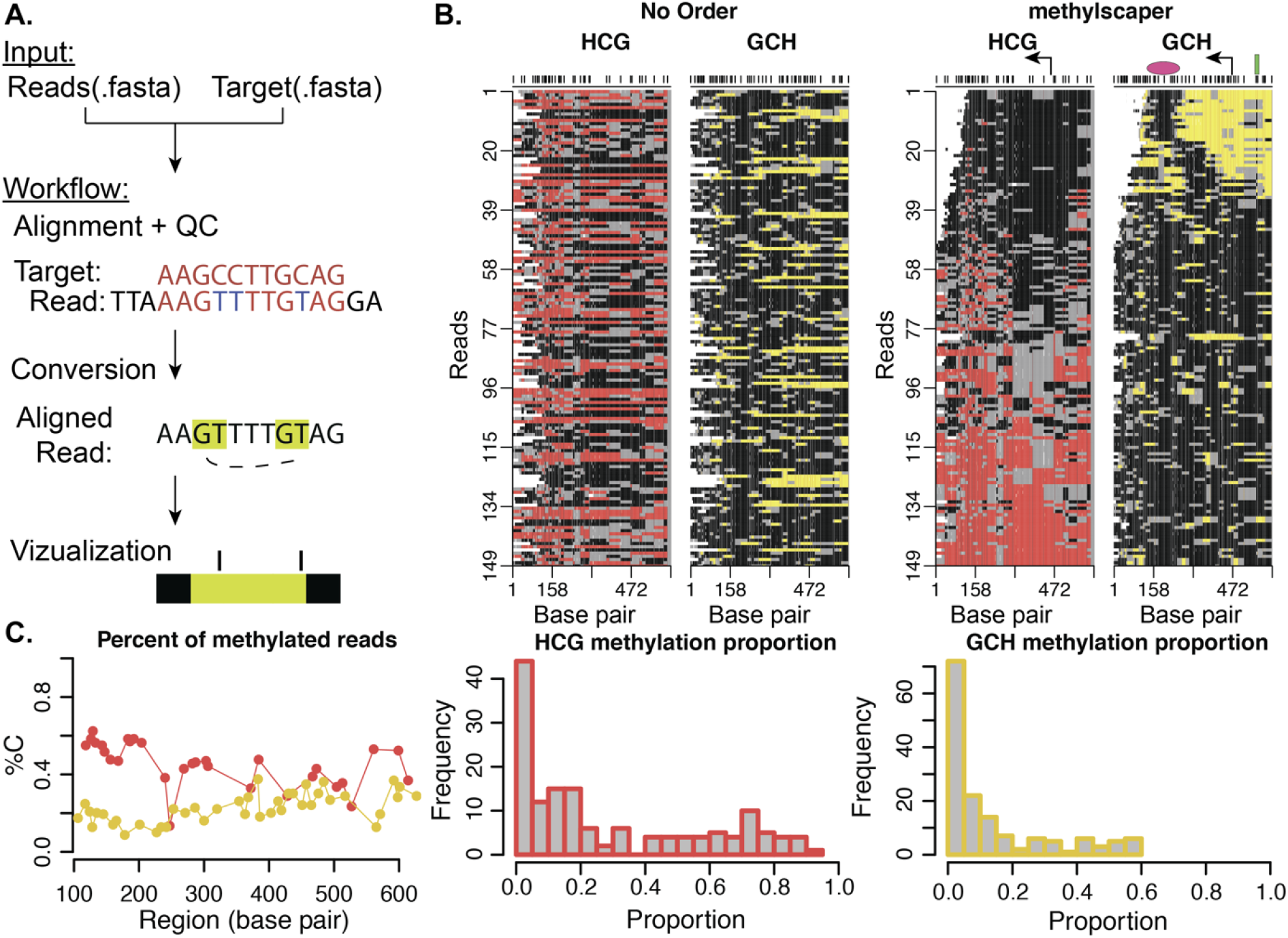
An overview of methylscaper. A. Flowchart of the bioinformatic pre-processing pipeline. B. methylscaper plots of the MAPit-BGS data, generated with two different orderings. The data in the left plot is not ordered; the data on the right was ordered with methylscaper’s weighted principal component algorithm. A pink oval was added to indicate the ~150-bp +1 nucleosome downstream of the transcription start site; a green rectangle was added to indicate a sequence-specific DNA-binding factor. C. Summary plots generated by methylscaper. Left: The percentage of reads methylated at each base pair. Center: A histogram of the proportion of HCG sites that are methylated in each read. Right: A histogram of the proportion of GCH sites that are methylated in each read.

Methylscaper then orders the single-molecule reads to visualize heterogeneity, allowing identification of patterns of endogenous methylation, as well as transcription factor and nucleosome occupancy. Using a numerical key to represent patches of methylation patterns, methylscaper constructs a matrix containing both endogenous (HCG) and induced (GCH) methylation states for the set of reads. Weighted-PCA is performed on the entire matrix, with reads assigned a weight based on the number of methylation patches between two fixed base pairs chosen by the user. This allows the weighting to focus on either type of methylation and to emphasize specific genomic regions (Supplementary Figure 1). The first weighted principal component is used to determine the global read order; as shown in Supplementary Figure 2, the first component is highly correlated with methylation and accessibility.

Following the determination of the global ordering, users can perform an optional second-stage refinement step, in which a contiguous subset of the reads is reordered using the PCA procedure to increase the resolution of patterns (Supplementary Figure 3). Additional statistics are also calculated from the reads that are then comparable across datasets or treatments. In Figure 1C, methylscaper calculates experiment-wide statistics, such as the proportion of methylated sites at each base across all reads, as well as read-specific statistics including the proportion of each read that is GCH or HCG methylated.

## Results

We applied methylscaper to a dataset with 149 single-molecule reads generated using MAPit-BGS. This dataset is from an epigenetic study of methylation resilencing in the *EMP2AIP1* promoter region following withdrawal of the DNA methyltransferase inhibitor 5-aza-2’-deoxycytidine using cell line RKO. A comparison of the visualization without any ordering versus our weighted PCA is shown in Figure 1B. Without any ordering of the read structure (left panel), drawing biological conclusions is precluded. Using methylscaper (right panel), it becomes evident that endogenous HCG methylation (red-gray) inversely correlates with GCH accessibility (yellow-gray). The quality of ordering also allows visualization of the +1 nucleosome sliding across cells–the ~150 bp footprints (black areas) that move in register with expansion/shortening of the accessible nucleosome-free region at the transcription start site (TSS). In reads 25-35, two phased nucleosomes are observed, punctuated by an accessible linker. Finally, protection of two GCH sites upstream of the TSS and within the nucleosome-free region detects binding of a sequence-specific transcription factor.

We also compared visualization with methylscaper to existing tools. In previous manuscripts using MAPit-BGS, hierarchical clustering alone was used to order the reads, but we have found this method fails with increasing complexity of patterns and number of reads and often breaks the reads into distinct blocks that have locally optimal orderings, but are out of order with respect to a global structure (Supplementary Figure 4). When patterns in the data are heterogeneous and many reads are available, this leads to unorganized and potentially uninformative visualizations. Line plots, also commonly used to visualize methylation and accessibility status (for example, as implemented in the aaRon R package(20) or the NOMePlot software), either present the status of a single read at a time or of a moving population average of statuses across all reads (Supplementary Figure 5). This type of plot is insufficient when visualizing a large number of reads, as using population averages often leads to a loss of critical information when methylation status or nucleosome occupancy is highly variable in heterogeneous cell populations. Commonly used lollipop plots also become unclear when a large number of reads are available (Supplementary Figure 5).

Next, we applied our results to a single-cell dataset generated using the scNMT-seq protocol that jointly profiles methylation and accessibility chromatin states in single cells(13). As shown in Clark et al., we also observe high levels of open chromatin near the TSS for *Eef1g*, though we find evidence of a +1 nucleosome approximately +250 bp downstream of the TSS (Supplementary Figure 6).

## Supporting information

Methylscaper Vignette

