## Supplemental Text and Figures for "methylscaper: an R/Shiny app for joint visualization of DNA methylation and nucleosome occupancy in single-molecule and single-cell data"

**Supplementary Methods**

1**. Description of methylscaper pre-processing procedure**
Methylscaper allows either single-molecule (e.g. MAPit-BGS) or single-cell (e.g. scNMT-seq) data to be given as input. Single-cell data should be initially processed with the Bismark software tool. We specifically use the output of the "bismark_methylation_extractor", filtered to the chromosome level (an example of this is given in the methylscaper package vignette).

Single-molecule sequence data should be given in the FASTA format, with a suitable reference file. The alignment is done using the pairwiseAlignment function in the Biostrings R package. We set the alignment penalty to 1 for exact matching and for C-T or G-A conversions and allow the alignment to occur either directly or to the reverse complement or both strands' reverse complement. Nonmatching alignments receive a penalty of -5. The gap opening penalty is -8. The highest scoring alignment is kept, and aligned reads are chosen if their score is higher than the maximal pairwise difference in scores. We do not include GCG sites because their status is biologically ambiguous.

After the preprocessing is complete, we assign all sites a numeric value. GCH sites that are methylated are assigned the value -4; unmethylated is assigned -1. Bases between two methylated GCH sites are assigned -3, and those between two unmethylated GCH sites are assigned -2. These in-between bases represent what we refer to as a methylation 'patch'. If two consecutive GCH sites do not have the same methylation state, the bases in between are assigned the value -2.5 resulting in a gray color. The same scheme with positive values is used for HCG sites. A gray patch may also be due to missing data, especially for sparse single-cell data in which the methylation status of a HCG or GCH site is unknown due to missed coverage in sequencing. This unique assignment provides a structure with which the data can be ordered using numerical methods and then visualized.

**2. Description of methylscaper ordering procedure**
The primary ordering method used by methylscaper is derived from a weighted principal components analysis. Let $X$ denote the $n$ x 2$b$ representational matrix, where $n$ denotes the number of reads (i.e. molecules or cells) and $b$ denotes the number of base pairs ($X$ is thus formed by joining the columns of the GCH and HCG matrices row-wise). The weight of row $i$, denoted $w_i$, is computed as the number of bases in methylated patches that lie within a region of the reads indicated by the user. Numerically, we compute $w_i$ *as*

$$w_i = \sum_{j=1}^{2b} I(X_{i,j})$$

where $I$ is an indicator function, equal to 1 if $X_{i,j}$ falls within a methylation patch of interest, and equal to 0 otherwise. We then normalize all of the weights so that they sum to 1, and multiply each row of $X$ by the square-root of its normalized weight, forming the weighted representational matrix $X^*$. We then compute the Singular Value Decomposition of $X^*$, i.e.

$$USV^T = X^*$$

The order of the entries in the first column of *U* is then used as the order of the reads in generating the methylscaper plots.

**3. Computational performance**
methylscaper's runtime performance was evaluated using the *microbenchmark* R package. We compared the principal component analysis (PCA) method with the hierarchical clustering (HC) method provided by the *seriation* package by running these methods on the MAPit-BGS data. To evaluate the performance of each method on larger scale data, we replicated the rows of the representational matrix 2, 5, and 10 times to increase the total number of reads. Methylscaper scales to analyzing large datasets while illuminating heterogeneous epigenetic features that will be useful as single-cell approaches continue to evolve. Summary statistics after 100 runs of each method are given below in Supplementary Table S1. All times are given in milliseconds.

**Supplementary Table S1**

| method | dataset | min | mean | max |
|---|---|---|---|---|
| HC | 1x | 217.776 | 232.5715 | 526.722 |
| HC | 2x | 423.127 | 444.74145 | 696.351 |
| HC | 5x | 1206.754 | 1243.1683 | 1588.338 |
| HC | 10x | 2897.589 | 3176.36745 | 3805.473 |
| PCA | 1x | 247.221 | 265.06005 | 549.325 |
| PCA | 2x | 474.103 | 492.76629 | 812.351 |
| PCA | 5x | 1322.156 | 1398.77479 | 1741.466 |
| PCA | 10x | 3182.073 | 3370.89101 | 3828.59 |

**4. Additional details on analysis of case-study datasets**
i) MAPit-BGS: The raw data is included in Supplementary Data. The file ref.Pacbio.fa is the sequence of the promoter region of the EMP2AIP1 gene. The file seq_file.fasta are the sequences of all reads generated by the MAPit-BGS experiment. After quality control in the bioinformatic pipeline there were 149 high quality single molecule reads. We ran methylscaper with weighting on the HCG features from position 308 to 475, we then iteratively refined reads from 1-54, 1-39, and 1-24.
ii) scNMT-seq: We downloaded the scNMT-seq data from GSE109262, which contained 61 cells that passed quality control in the original publication in Clark et al. 2018. We focused on a region of the *EEFG1* gene from (TSS-200, TSS+500), where the TSS is located on chromosome 19 at 8,967,041 bp. We ran methylscaper with weight on the GCH features from base 47 to 358 and refined reads from 27 to 42.

### Supplementary Figures

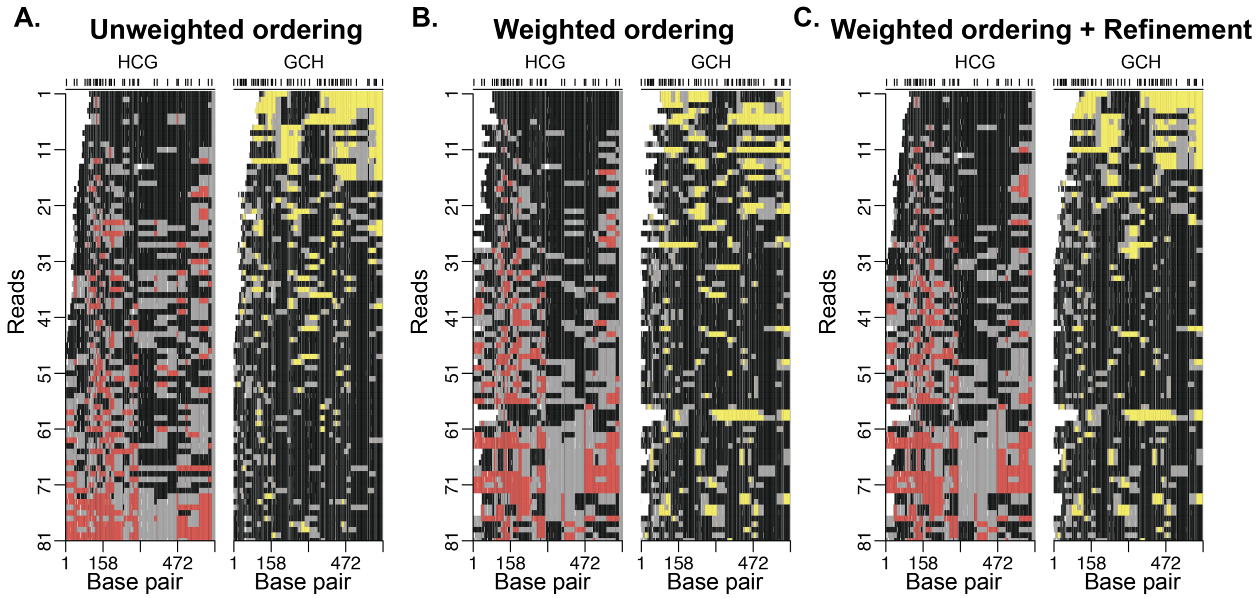


Supplementary Figure 1: A comparison of ordering procedures provided by methylscaper using reads 20-100 from Figure 1 as an example. A. methylscaper plot ordered using the unweighted PCA. B. methylscaper plot of data ordering with a weighted PCA where ordering is focused on the red patch from 308bp to 475bp. C. The weighted methylscaper plot with weighting as in (B) and with refinement ordering on the first 40 molecules. Now, both the HCG and GCH methylation patches are clearly visible.

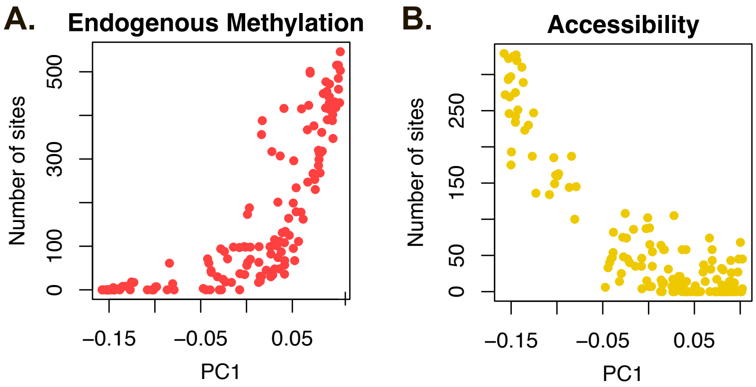


Supplementary Figure 2: Scatter plots of the number of methylated sites versus the first principal component of the representational state matrix. A. Number of endogenously methylated sites versus PC1 values. B. Number of accessible sites versus PC1 values.

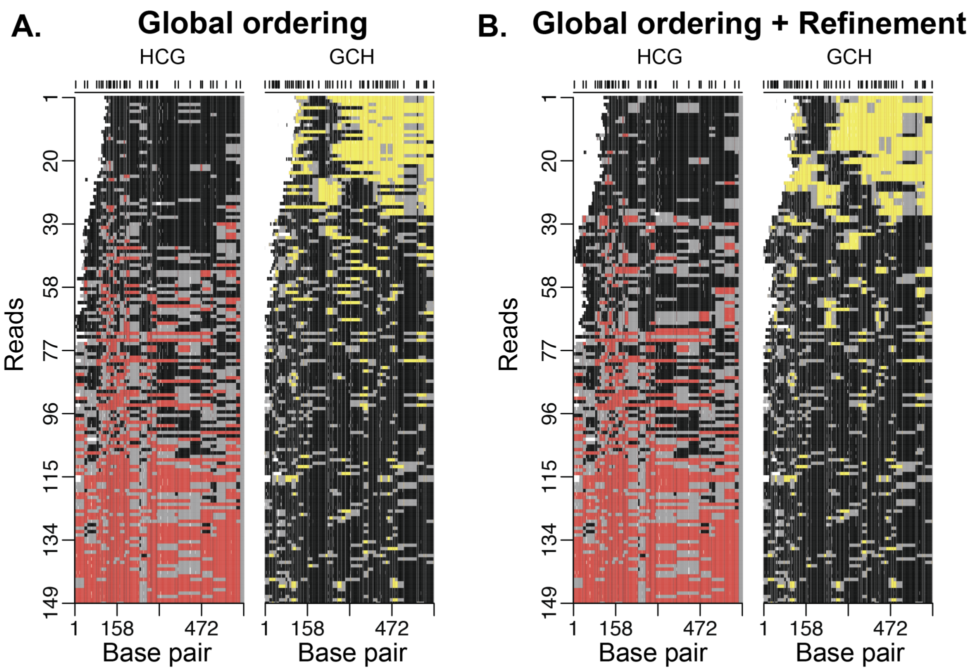


Supplementary Figure 3: A comparison of unrefined and refined methylscaper plots. A. methylscaper plot after the initial global ordering by PCA. B. The methylscaper plot after refining the ordering of a subset of the first 70 reads, followed by refinement of the first 35 reads.

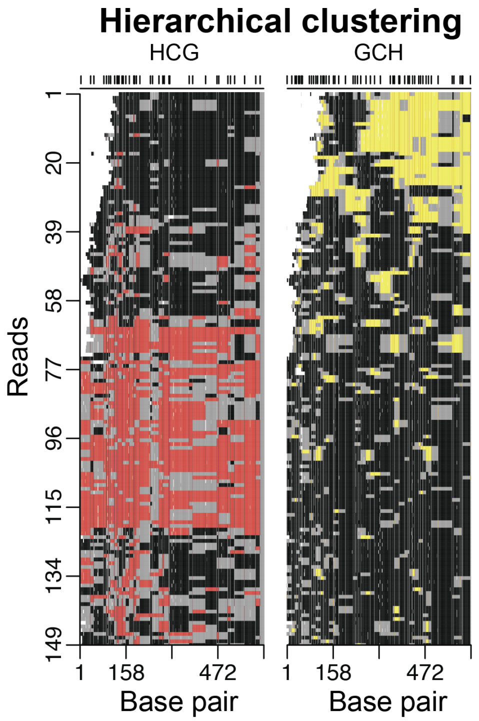


Supplementary Figure 4: A methylscaper plot with the read ordering computed with Hierarchical Clustering.

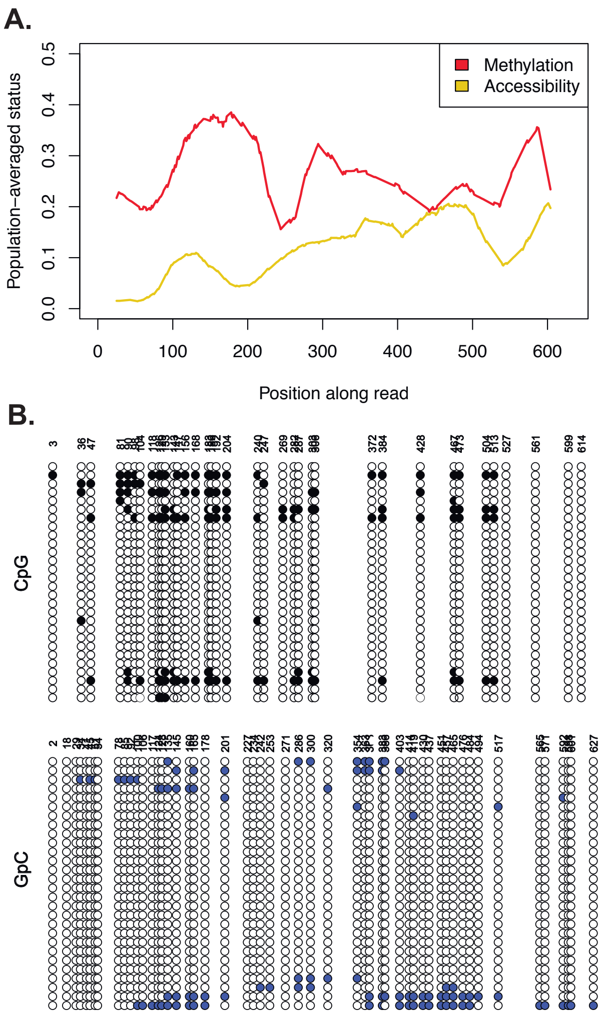


Supplementary Figure 5. Examples of alternative plots used to visualize methylation data. (A) A line plot of the moving average (50bp window) of methylation and accessibility across all reads. (B) A Lollipop plot generated from the NOMePlot software with a subset of 28 reads.

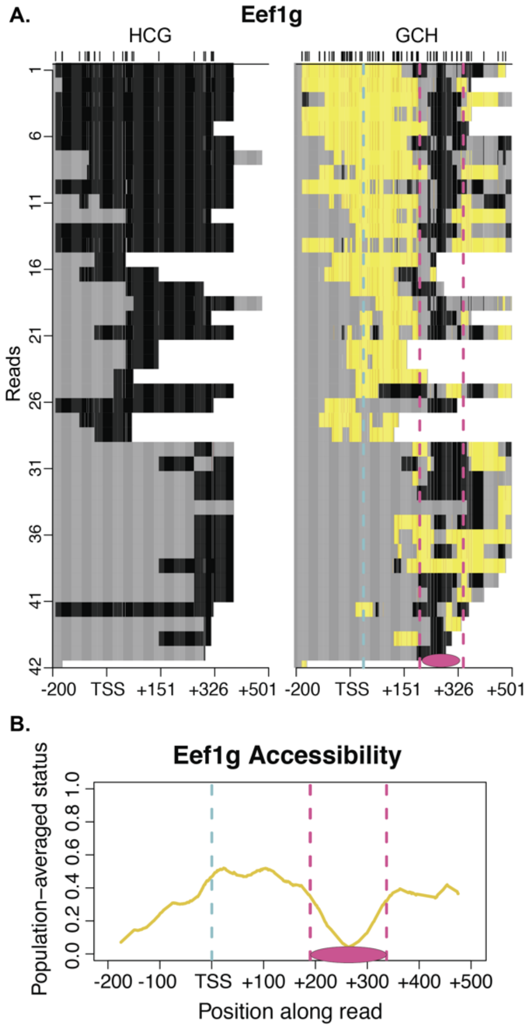


Supplementary Figure 6: Analysis of the Eef1g gene from a scNMT-seq dataset. A. Methylscaper plot of Eef1g surrounding the transcription start site (TSS). The region immediately near (+/- 200 bp) the TSS is highly accessible, while a nucleosome downstream of the TSS is centered around +250bp. B. A moving average (50bp window) of accessibility across all reads also indicates evidence of a nucleosome.
