## Supplementary material for "methylscaper: an R/Shiny app for joint visualization of DNA methylation and nucleosome occupancy in single-molecule and single-cell data": Methylscaper Vignette

### Introduction

methylscaper is an R package that provides functions for processing and visualizing data generated by methods jointly profiling methylation and chromatin accessibility (MAPit, NOMe-seq, scNMT-seq, nanoNOMe, etc.). The package offers processing for both single-cell and single-molecule data, and a common interface for jointly visualizing both data types through the generation of ordered representational methylation-state matrices. Users may also run the package through a Shiny app which provides an interactive seriation process of refinement and re-weighting that optimally orders the single DNA molecules to discover methylation patterns and nucleosome positioning.

#### Installation

Currently, the only way to install methylscaper is via the development version on GitHub.

```
install.packages(devtools)
devtools::install_github("rhondabacher/methylscaper", build_vignettes=TRUE)
library(methylscaper)
```

### Usage

#### Data processing

##### Single-cell data

methylscaper can handle the analysis of single-cell data from methods such as scNMT-seq processed with the Bismarck software tool. Specifically, we use the read-level output of the "bismarck_methylation_extractor" script to generate the methylation-state matrix which is used in further analysis.

Most of the processing is handled by the prepSC() function, but the input data does need to be filtered to the chromosome level. The code below demonstrates this using data from Clark et al., 2018, obtained from GSE109262. For the sake of this example, we assume that the GSE109262_RAW.tar directory is downloaded to ~/Downloads/.

```
setwd("~/Downloads/GSE109262_RAW/")
```

```r
cgfiles <- sort(grep("met", list.files("~/Downloads/GSE109262_RAW/"), value = T))
gcfiles <- sort(grep("acc", list.files("~/Downloads/GSE109262_RAW/"), value = T))

useChr <- "8" # we only save the input data for chromosome 8

cg.seq <- list()
for(i in 1:length(cgfiles)) {
  cg.seq[[i]] <- read.table(cgfiles[i], header=T, stringsAsFactors = F)
}

gc.seq <- list()
for(i in 1:length(gcfiles)) {
  gc.seq[[i]] <- read.table(gcfiles[i], header=F, stringsAsFactors = F,
                            colClasses = c("character", "numeric", "numeric"))
  colnames(gc.seq[[i]]) <- c("chr", "pos", "rate")
}

cg.seq.sub <- lapply(cg.seq, function(x) {
  QQ <- x[order(x$pos),]
  QQ = subset(QQ, chr==useChr)
  return(QQ)
})


gc.seq.sub <- lapply(gc.seq, function(x) {
  QQ <- x[order(x$pos),]
  QQ = subset(QQ, chr==useChr)
  return(QQ)
})
```

Now that the data is filtered to the chromosome level, we can use `methylscaper` to generate the methylation-state matrix. The `prepSC` function requires users to indicate a subset of the reads to process, via the `startPos` and `endPos` arguments. This subset generally corresponds to a gene of interest; for this example, we use the Ctcf gene.

```r
prepsc.out <- prepSC(gc.seq.sub, cg.seq.sub, startPos = 105636488, endPos = 105636993)
gch <- prepsc.out$gch
hcg <- prepsc.out$hcg
```

The `gch` and `hcg` objects are matrices representing accessibility and methylation status, respectively, and are used by other `methylscaper` functions for visualization.

#### Single-molecule data

`methylscaper` can process single-molecule data from .fasta files (given an appropriate reference file) with the `runAlign` function. The sequences are aligned to the reference with the `Biostrings` package, and then mapped to the methylation- and accessibility-state matrices. For very large datasets, the align function may be run on high-throughput servers rather than locally.

```
fasta <- read.fasta("seq_file.fasta")
ref <- read.fasta("ref.Pacbio.fa")
align.out <- runAlign(fasta = fasta, ref = ref)
gch <- align.out$gch
hcg <- align.out$hcg
```

#### Seriation and visualization

Once the methylation- and accessibility-state matrices have been generated, we can can generate an ordering of the reads and visualize them with a sequence plot. To demonstrate this, we use an example dataset included in the package called `day7`.

The `initialOrder` function computes an ordering of the state matrices, using a given method. By default, the function uses a PCA-based ordering which we find optimal and efficiencly scales to large datasets, but technically any method supported by the `seriation` package can be used. It can also perform a weighted ordering, according to the methylation or accessibility status of a subset of the data's columns.

```
data(day7)
gch <- day7$gch
hcg <- day7$hcg
orderObj <- initialOrder(gch, hcg, Method = "PCA")
```

`initialOrder` returns an object of class `orderObject`, which includes the combined state data matrix and the ordering of the reads. We then generate a sequence plot with the `plotSequence` function.

```
plotSequence(orderObj)
```

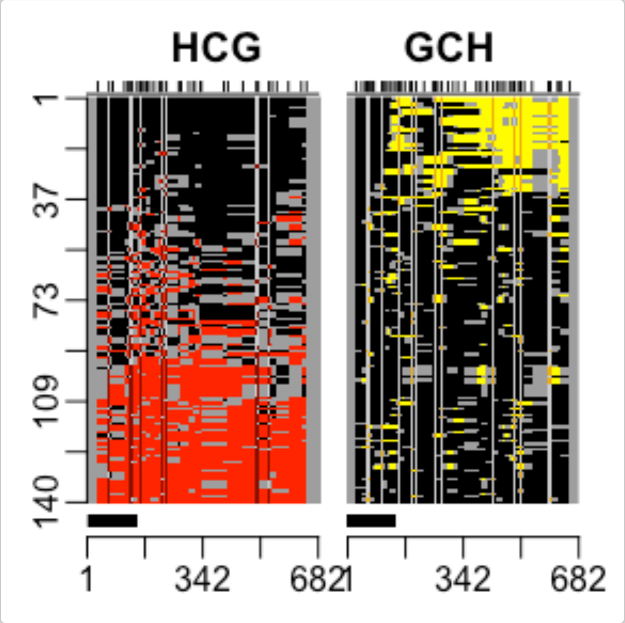


The plot on the left corresponds to methylation status, and the right to accessibility status The red and yellow colored portions represent the areas on each read that are methylated and accessible, respectively.

We can also refine the ordering of the reads with `refineFunction`, which reorders a subset of the reads with a given method. The code below reorders the first 50 reads and generates a new sequence plot.

```
orderObj$order1 <- refineFunction(orderObj, 1, 50)
plotSequence(orderObj)
```

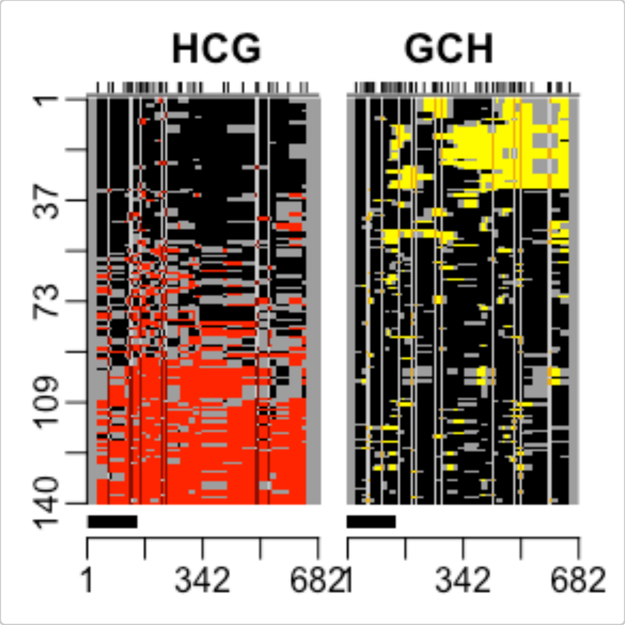


We can also generate basic summary plots of the data.

```
percent.c.out <- percent_C(orderObj, plotPercents = TRUE)
```

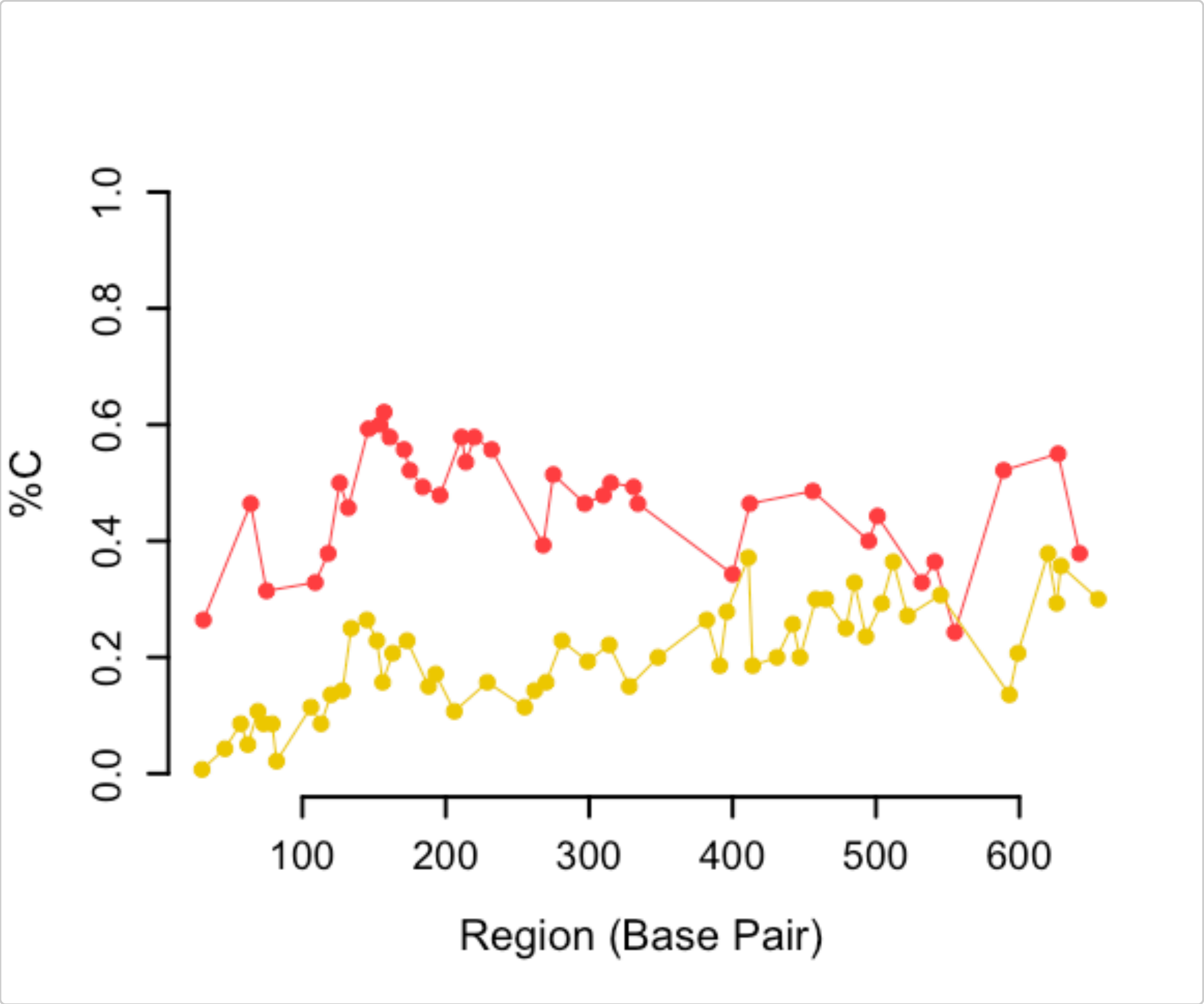


```
prop.color.out <- proportion_color(orderObj, color = "YELLOW", plotHistogram = TRUE)
```

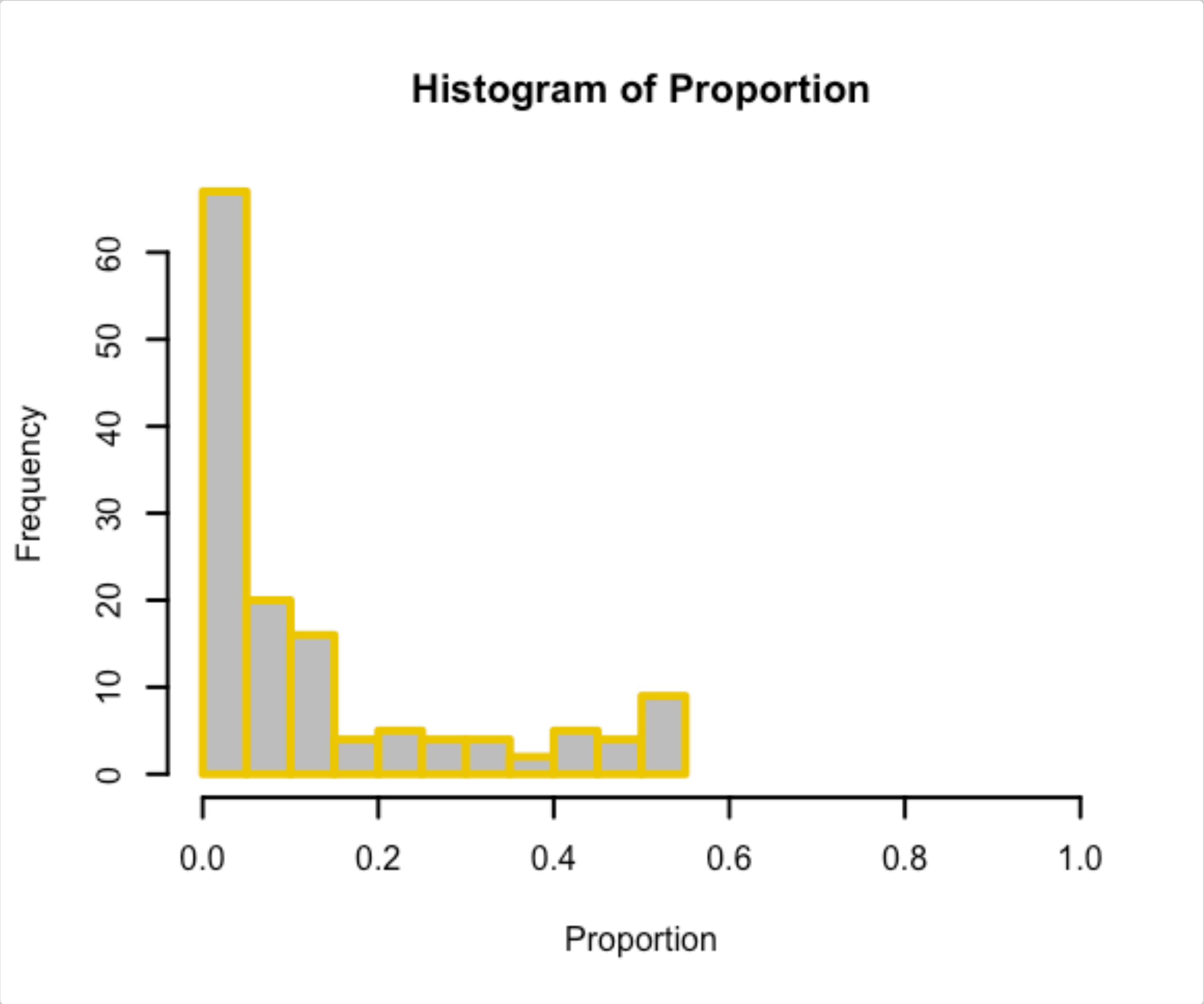


#### Shiny App

The Shiny app offers all of the package functionality described above, and can be run by calling the function `methylscaper()` from an R session with the package loaded. The app is organized into single-cell and single-molecule panels, which each have tabs for processing and visualization.

In the "Single-cell" panel, users upload the raw data (after being filtered to the chromosome level) as .rds files and then enter start and end position values to initiate the processing. After the initial processing is complete, the Shiny app generates a slider that can be used to adjust the selected region.

Once the initial sequence plot is generated, `methylscaper` allows the user to dynamically refine and re-weight the plot via Shiny's brushing mechanics. Clicking and dragging along either of the two plots will select sites (i.e., columns) by which to weight the data in the site matrix. `methylscaper` will then regenerate the plot with a new ordering, influenced by the weighted sites. With "PCA" selected as the seriation method, the new ordering will be generated with a weighted Principal Components Analysis. If "ARSA" is selected, the ordering is found by first building a weighted Euclidean distance matrix, which is then passed to the Simulated Annealing algorithm. Note that weighting is done with respect to either the GC sites or the CG sites - the plot on which brushing is performed determines which sites to use.

If "Refinement" is selected as the brushing option, clicking and dragging on the sequence plot will select reads (i.e., rows) to reorder locally. PCA is used by default, but Hierarchical Clustering can also be selected as the refinement method. Unlike re-weightings, refinements to the sequence plot stack onto each other, and several refinements can be done to a single plot before exporting. However, it is important to note that re-weighting the sites will reorder the entire set of data, and hence will undo any refinements that you may have made.

The green lines indicate the columns selected for weighting, and the blue lines indicate the rows that were most recently refined.

After making any desired changes, the sequence plot can be saved as either a PNG, PDF, or SVG file. Additionally, `methylscaper` keeps track of all changes made to the plots in the form of a changes log, which can be saved as a text file.

The “Single-molecule” panel includes a “Preprocessing” tab, where users are prompted to input two sequence FASTA files, the first containing all of the raw reads and second containing the reference sequence for the region of interest. Users are asked to enter two files paths, indicating where to save the two processed state matrices. The GCH file contains the GC site accessibility data, and the HCG file contains CG site methylation data. These files can then be loaded on the “Seriation” tab, which offers the same functionality as described above.
